# Genomic surveillance of *Escherichia coli* ST131 identifies local expansion and serial replacement of subclones

**DOI:** 10.1101/814731

**Authors:** Catherine Ludden, Arun Gonzales Decano, Dorota Jamrozy, Derek Pickard, Dearbhaile Morris, Julian Parkhill, Sharon J. Peacock, Martin Cormican, Tim Downing

## Abstract

*Escherichia coli* sequence type 131 (ST131) is a pandemic clone that is evolving rapidly with increasing levels of antimicrobial resistance. Here, we investigated an outbreak of *E. coli* ST131 producing extended spectrum β-lactamases (ESBLs) in a long-term care facility (LTCF) in Ireland by combining data from this LTCF (n=69) with other Irish (n=35) and global (n=690) ST131 genomes to reconstruct the evolutionary history and understand changes in population structure and genome architecture over time. This required a combination of short and long-read genome sequencing, *de novo* assembly, read mapping, ESBL gene screening, plasmid alignment and temporal phylogenetics. We found that clade C was the most prevalent (686 out of 794 isolates, 86%) of the three major ST131 clades circulating worldwide (A, B, C), and was associated with the presence of different ESBL alleles, diverse plasmids and transposable elements. Clade C was estimated to have emerged in ∼1985 and subsequently acquired different ESBL gene variants (*bla*_CTX-M-14_ vs *bla*_CTX-M-15_). An ISEcp*1-* mediated transposition of the *bla*_CTX-M-15_ gene further increased the diversity within Clade C. We discovered a local clonal expansion of a rare C2 lineage (C2_8) with a chromosomal insertion of *bla*_CTX-M-15_ at the *mppA* gene. This was acquired from an IncFIA plasmid. The C2_8 lineage clonally expanded in the Irish LTCF from 2006, displacing the existing C1 strain (C1_10) and highlighting the potential for novel ESBL-producing ST131 with a distinct genetic profile to cause outbreaks strongly associated with specific healthcare environments.

**Importance:** Extraintestinal pathogenic *E. coli* (ExPEC) ST131 is adapting in the context of antibiotic exposure, resulting in a pandemic with distinct genetic subtypes. Here, we track the evolution of antibiotic-resistance gene variants originally discovered in an ExPEC ST131 outbreak that was identified in a LTCF in Ireland. Analysis of 794 global ST131 genomes show that subclade C1 was associated with the initial infection outbreak, but that a new lineage from subclade C2 successfully displaced C1. This genetically distinct C2 subclade with a chromosomal insertion of a key antibiotic-resistance gene had clonally expanded within the LTCF. We provide new insights into the timing of genetic events driving the diversification of C2 subclades to show that that outbreak C2 strain likely evolved elsewhere before spreading to the LTCF. This study highlights the scope of antibiotic-resistance gene rearrangement within ST131, reinforcing the need to integrate genomic, epidemiological and microbiological approaches to understand ST131 transmission.

## Introduction

*Escherichia coli* is the leading cause of urinary tract infections and bloodstream infections (BSIs) (1, 2), with the number of *E. coli* BSIs continuing to increase in Europe and the United States since the early 2000s (3–7). This has been associated with the emergence and dissemination of antibiotic-resistant *E. coli* producing extended-spectrum β-lactamases (ESBL-*E. coli*) conferring resistance to many beta-lactam antibiotics, including cephalosporins (6, 7). Infections caused by ESBL-*E. coli* are associated with higher morbidity and mortality, longer hospital stays and higher healthcare costs compared to infections with antibiotic-susceptible *E. coli* (1, 8–10).

The global spread of ESBL-*E. coli* is largely attributed to the dissemination of *E. coli* strains carrying the *bla*_CTX-M-15_ gene, especially *E. coli* O25b:H4-ST131. Genomic analyses estimated that ST131 emerged in North America over 30 years ago, coinciding with the first use of fluoroquinolone (FQ) in 1986 (11, 12). Previously, three major lineages of ST131 were identified that differed mainly based on their *fimH* alleles: A (mainly *fimH*41), B (mainly *fimH*22) and C (mainly *fimH*30) (13). Clade C has predominated since the 2000s, corresponding with the rapid dissemination of the *bla*_CTX-M-15_ allele (13, 14). Clade B also contains the subclade B0 which differs phylogenetically from the remaining B isolates by carrying *fimH*27 and is considered ancestral to Clade C (13, 15). Clade C consists of three subclades termed C0, C1 and C2. Clade C0 has been reported as ancestral and is composed of FQ-susceptible isolates. In contrast, clades C1 (also known as *H30*R) and C2 (also known as *H30*Rx) are characterised by a double mutation at the *gyrA* and *parC* genes conferring high-level resistance to FQ (11, 13, 16). Clade C2 is sub-divided from C1 based on specific SNPs at *fimH*30 as previously described and is associated with the *bla*_CTX-M-15_ gene (16, 17).

ST131 has principally been associated with the hospital setting, though in recent years it has also been reported at high prevalence in the community (18–20). There is increasing evidence that ST131 is common in the elderly and that long-term care facilities (LTCFs) are important reservoirs for ESBL-producing ST131. Reported rates of multidrug-resistant (MDR) *E. coli* ST131 carriage in residents of LTCFs include 55% in Ireland, 36% in the UK and 24% in the United States (21–23). It is projected that the proportion of the European Union population aged ≥ 65 years and ≥ 80 years will increase to 29% and 11.5 % by 2060, respectively (24). This will likely lead to a rise in the number of people residing in LTCFs, potentially expanding the reservoir of ESBL-producing ST131. Infection control measures targeting *E. coli* have focused primarily on hospitals, and there is still a limited understanding of *E. coli* transmission dynamics within LTCFs, and between hospitals and LTCFs (22, 25). To develop effective strategies for containment and prevention of infections, it is necessary to improve our ability to detect transmission events and to monitor the emergence of new clones. Here, we used short and long read genome sequencing to investigate an ESBL-*E. coli* ST131 outbreak in a LTCF in Ireland. We describe the genetic basis of antibiotic resistance and the evolution of ESBL-*E. coli* ST131 over a seven-year period. We focused our analyses on ST131 clade C because of its high frequency in this LTCF, and its MDR profile. We analysed the population structure and inferred the evolutionary history of the LTCF isolates in the context of a local hospital and global collections of *E. coli* ST131 to further our understanding of its epidemiology.

## Results

### ESBL gene profiles among an *E. coli* ST131 outbreak in Ireland

In this study, we focused on the genetic profiles of 90 *E. coli* ST131 (local collection) isolated between 2005 and 2011 in Ireland, of which 69 were from one LTCF where an outbreak of ESBL-*E. coli* was first detected in 2006 (26). The other isolates were from other LTCFs (n=9), the referral hospital (n=10) and the community (n=2) (Supplementary Table 1). Initial screening of the 90 isolates indicated that 64 were *bla_CTX-M-15_*-positive, 17 were *bla_CTX-M-14_*-positive, one was *bla_CTX-M-27_*-positive, and four were positive for both *bla_CTX-M-15_* and *bla_CTX-M-14_* (Supplementary Table 1). Resistance to meropenem and ertapenem was not detected. Ribosomal sequence typing (rST) demonstrated a high incidence of rST1850 (44/90, 49%) (Supplementary Table 1), suggesting emergence of a unique local epidemic clone.

### ST131 clade C predominates in Ireland and elsewhere

We analysed the 90 isolates from the local collection in the context of a global collection of 704 *E. coli* ST131 genomes that contained four additional isolates from the referral hospital described in the local collection and 10 isolates from other hospitals in Ireland. To better understand the global population structure of *E. coli* ST131, we reconstructed the phylogeny of all 794 isolates based on a core genome alignment containing 12,518 SNPs (Figure 1). This recapitulated the three established ST131 clades (A, B and C) (29) and showed that most isolates were from C (n=686, 86.4%) followed by B (n=75, 9.4%) and A (n=33, 4.2%). The clade classification was supported by previously described *fimH* allelic differences (16): clade A was largely *fimH41* (30 out of 33), clade B *fimH22* (60 out of 70), subclade B0 *fimH27* (4 out of 5) and clade C *fimH30* (679 out of 686) (Table 1).

**Figure 1.**
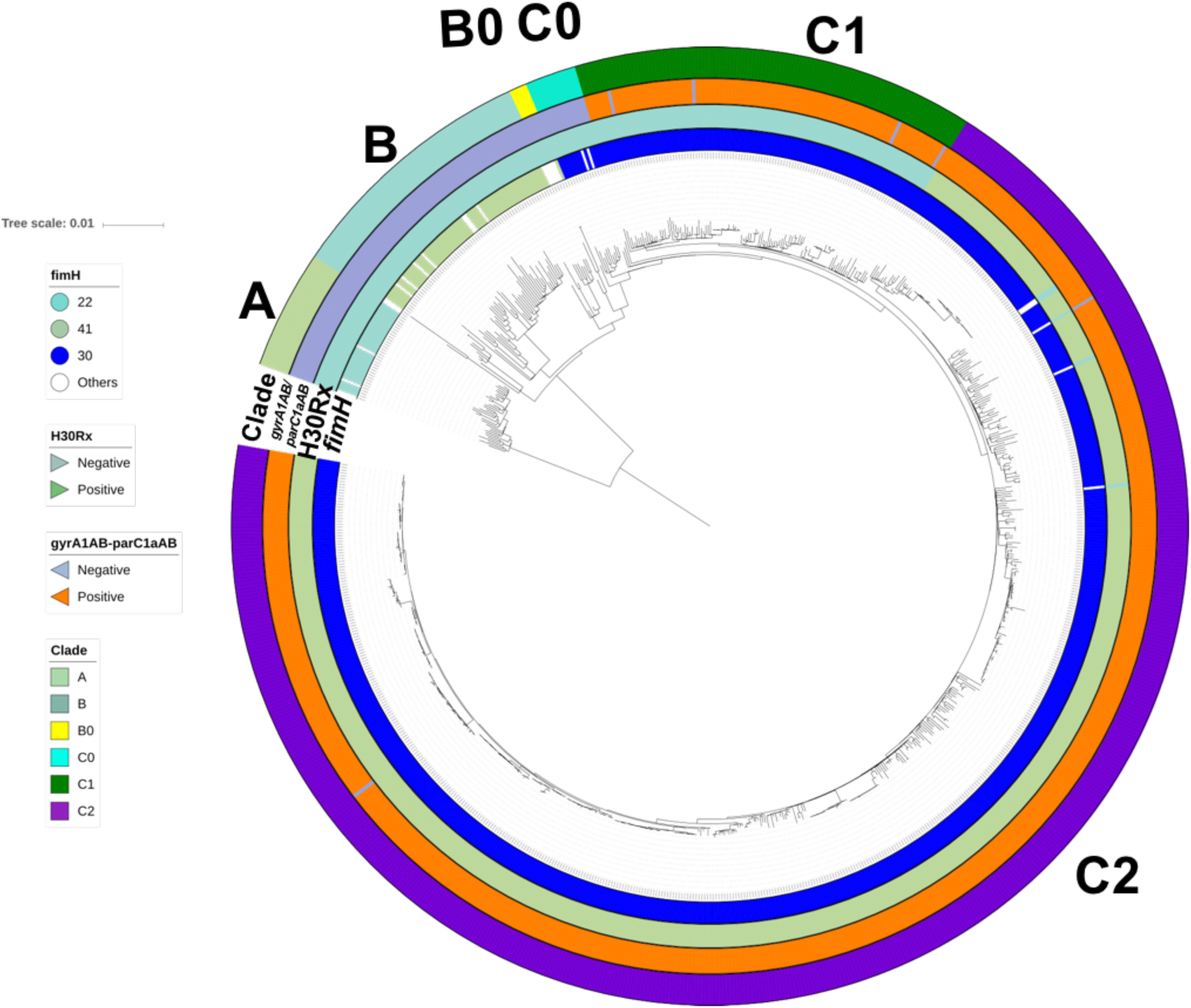
Phylogenetic reconstruction of N=794 global ST131 strains. Maximum likelihood phylogeny of n=794 global ST131 showed three main clades A (n=33), B (n=70), B0 (n=5) and C (n=686) with three common subclades in C: C0 (n=14), C1 (n=111) and C2 (n=561). The mid-point rooted phylogram was constructed with RAxML from the chromosome-wide SNPs arising by mutation, and visualized with iTol. Allelic profiling of fimH, gyrA-parC, the H30Rx phenotype, and clade classification are represented in colored strips around the phylogenetic tree.

**Table 1.**
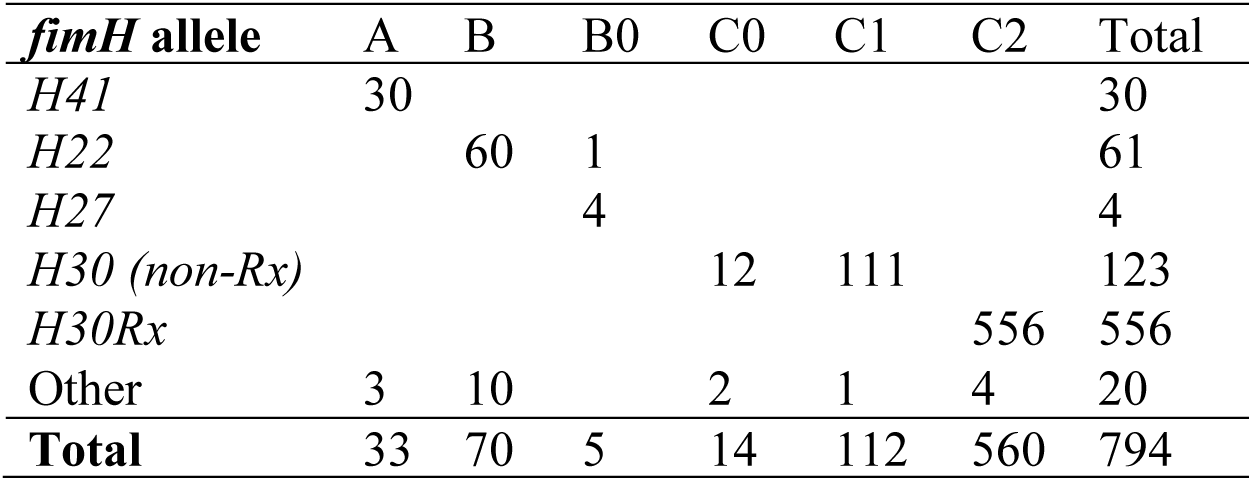
The entire ST131 set (n=794) consisted of three main clades sub-divided into six subclades: A (n=33), B (n=70), B0 (n=5), C0 (n=14), C1 (n=111) and C2 (n=561). The frequencies of the four most common *fimH* allele types are shown: *H41*, *H22*, *H27* and *H30*; the rest are listed as “other”. No FQ-resistance mutations were detected in *fimH22/27/41*.

FQ-resistance alleles *gyrA1AB* and *parC1aAB* (27) were present in nearly all C1 (96%) and C2 (99.7%) isolates, along with the *fimH30* allele, contrasting with their absence from the clades A, B, and B0 (Supplementary Table 1). This indicated that the Clade C ancestor acquired the *fimH30* allele and then differentiated into FQ-S (*H30*S or C0) and FQ-R (*H30*R or C1, *H30*Rx or C2) subclades. A limited number of C1 (n=1) and C2 (n=4) isolates had lost the FQ-R *gyrA1AB*-*parC1aAB* genotype, consistent with intermittent recombination at these and the *fimH* genes (11).

Considerable diversity within Clade C was demonstrated by the genetic clusters identified by Fastbaps (Figure 2 and Table 2): C0 (n=14, Fastbaps clusters 2-5 and 11), C1 (n=111, Fastbaps cluster 10), and C2 (n=560, Fastbaps clusters 7-9). All 104 Irish ST131 from the National Collection (local = 90, additional Irish isolates = 14, see Methods) were from clade C and there were no major differences in the rates of C0, C1 and C2 in the National collection (1%, 23% and 75%, respectively) compared to the global isolate collection (2%, 12%, 70%, respectively).

**Figure 2.**
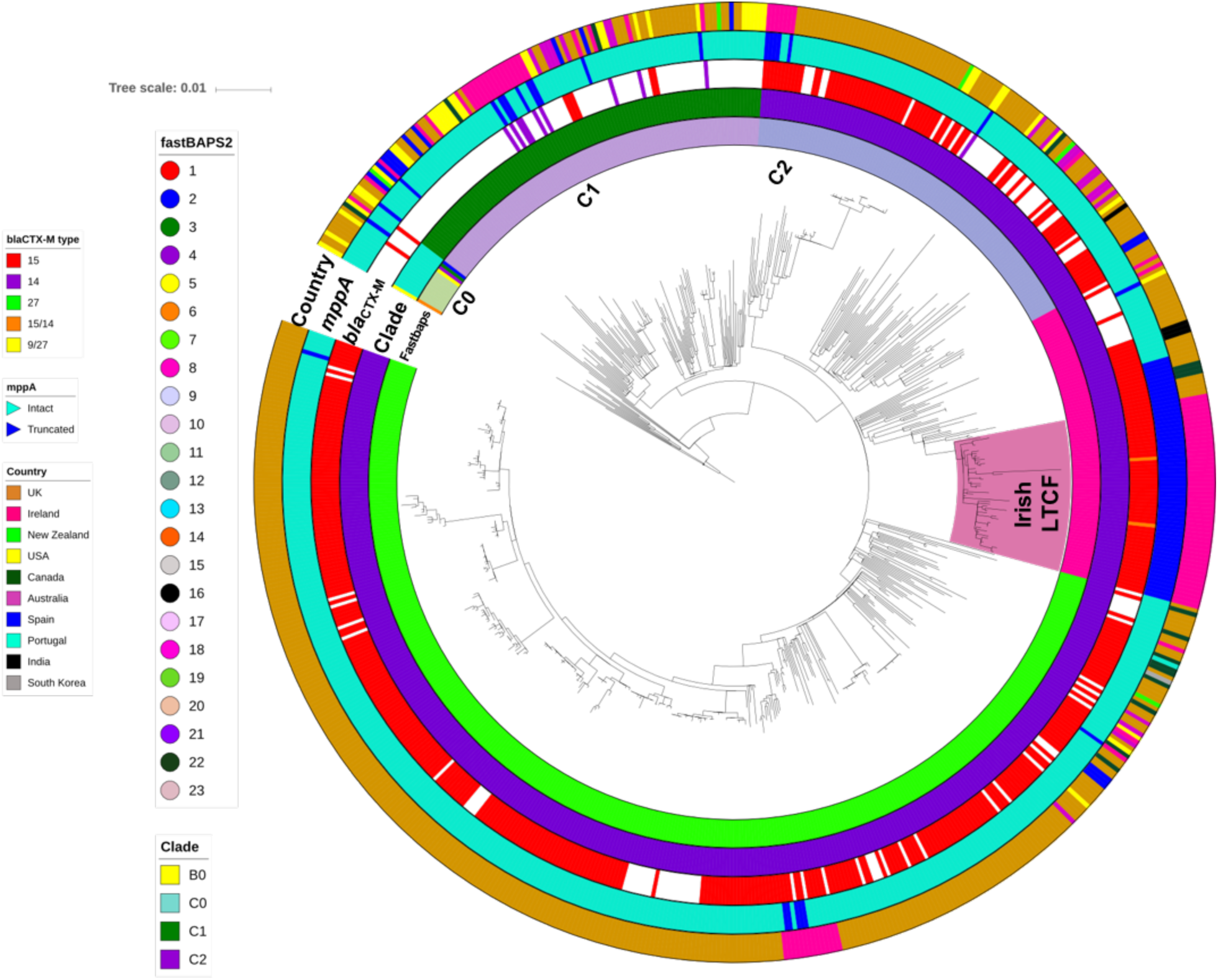
Maximum likelihood phylogeny of Clade C strains from the global ST131 collection. Phylogenetic reconstruction of 686 strains from Clade C with B0 as the outgroup. This shows 3 common subclades in C: C0 (n=14), C1 (n=111) and C2 (n=561) where the latter had three distinct subgroups: C2_7 (n=362, Fastbaps cluster 7), C2_8 (n=86, Fastbaps cluster 8) and C2_9 (n=113, Fastbaps cluster 9). Colored strips surrounding the phylogram represent the clade classification, Fastbaps clusters, *bla_CTX-M_* allelic profile, *mppA* state (intact or truncated) and the country of origin of each strain. The highlighted “Irish LTCF” clade was in C2_8.

**Table 2.** The entire ST131 set (n=794) was largely composed of isolates from clade C (n=686, 86% of total) that was categorised into five subclades by Fastbaps clustering: C0 (n=14, clusters 2-5 and 11), C1 (n=111, cluster 10), C2_7 (n=362, cluster 7), C2_8 (n=86, cluster 8) and C2_9 (n=113, cluster 9). The National (n=104) and global (n=690) ST131 had two main ribosomal sequence types (rSTs): rST1850 associated with the Irish C2_8 LTCF set (85%), and rST1503 that often corresponded to C2_7 (92.5%). Fastbaps clusters 2, 3, 4 and 5 in C0 represented one isolate each – only cluster 3 was *bla_CTX_*_-M-15_-positive.

|  | **Subclade**<br>**Fastbaps** | **C0**<br>**2-5,11** | **C1**<br>**10** | **C2_7**<br>**7** | **C2_8**<br>**8** | **C2_9**<br>**9** | **Totals** |
| --- | --- | --- | --- | --- | --- | --- | --- |
| **National**<br>**collection** | rST1503 | 1 | 24 | 16 | 1 | 7 | 49 |
|  | rST1850 |  |  |  | 45 |  | 45 |
|  | Other rSTs |  | 1 | 1 | 7 | 1 | 10 |
|  | Total | 1 | 25 | 17 | 53 | 8 | 104 |
| **Global**<br>**collection** | rST1503 | 11 | 82 | 319 | 21 | 101 | 535 |
|  | rST1850 |  |  |  | 11 |  | 11 |
|  | Other rSTs | 2 | 4 | 26 | 1 | 4 | 37 |
|  | Total | 13 | 86 | 345 | 33 | 105 | 582 |
| **Totals** |  | 14 | 111 | 362 | 86 | 113 | 686 |

### Phylogenetic reconstruction of three genetically distinct ST131 subclade C2 groups

Subclade C2 was structured into three Fastbaps clusters: 7 (n=362, named C2_7), 8 (n=86, C2_8) and 9 (n=113, C2_9) (Figure 2, Table 2). Most of the isolates in the National Collection (n=104) were represented by C2_8 (n=53, 51%), followed by C2_7 (n=17, 16%) and C2_9 (n=8, 8%). Within the global collection most isolates were C2_7 (n=345, 50%), with less in C2_8 (n=33, 5%) and C2_9 (n=105, 15%) (Figure 3). This showed C2_7 was more common globally than in Ireland (odds ratio = 3.1, p<6.5×10^-6^), and C2_8 was more widespread in Ireland than elsewhere (odds ratio = 10.6, p<2.2×10^-16^) (Figure 3).

**Figure 3.**
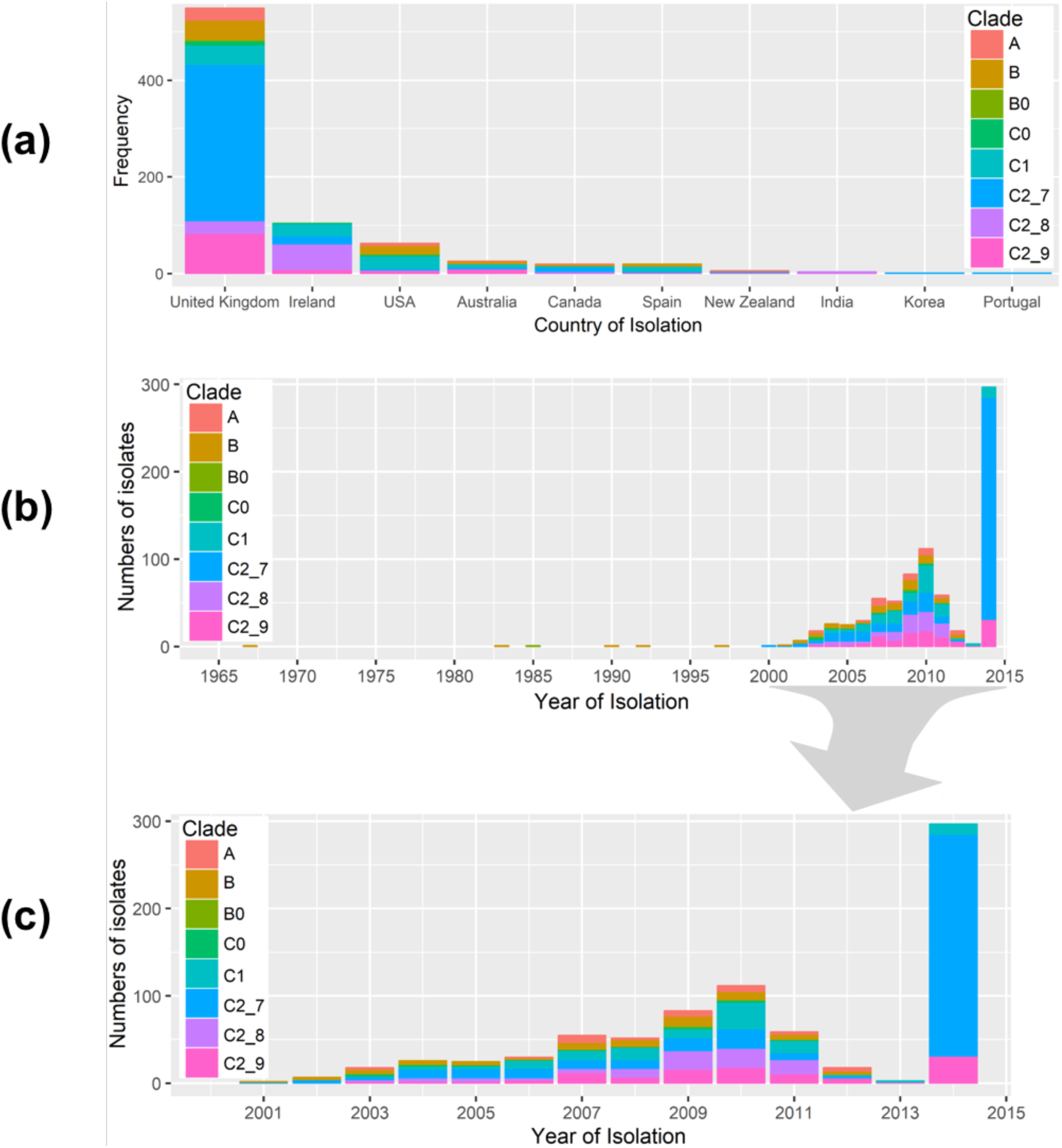
Geographic and temporal distribution of global ST131 isolates. ST131 from the eight subclades (n=794) showed differing frequencies across country of origin (a) and year of isolation (b and c). The subclades were A (n=33), B (n=70), B0 (n=5), C0 (n=14), C1 (n=111), C2_7 (n=362), C2_8 (n=86) and C2_9 (n=113). The ST131 were sampled during 1967-2014. The figures were generated using the ggplot2 and ggjoy packages in R v.3.5.2.

This difference was paralleled by the rST results, which showed that rST1503 was highly predictive of C2_7 globally (319 out of 345, 92.5%) and in Ireland (16 out of 17, 94%). Similarly, rST1850 was highly associated with C2_8 in Ireland (n=45, 85%), but less so for the global collection (11 out of 33, 33%, Table 2). This limited resolution suggests rMLST (ribosomal Multilocus Sequence Typing) has insufficient discrimination to accurately reflect the evolutionary history of clonal pathogens like ST131, and that core genome analysis was more informative.

### Long read sequencing uncovers chromosomal transposition of *bla_CTX-M_* genes

Five isolates from the Irish collection were selected for long-read sequencing to more accurately determine the location and genomic environment of the *bla_CTX-M-14_* and *bla_CTX-M-15_* genes. Four of five samples selected were *bla_CTX-M-15_* positive and members of Clade C2, of which three belonged to the predominant LTCF subclade (C2_8) and one from the predominant global clade (C2_7). The remaining long-read sequenced isolate was from Clade C1 and was *bla_CTX-M-14_*-positive. Each of the PacBio assemblies were used as references for Illumina read mapping for the collection of 794 isolates (see Methods). The three C2_8 PacBio genomes (ERR191646, ERR191657, ERR191663) demonstrated chromosomal insertion of a 2,971 bp ISEcp*1*-*bla_CTX-M-15_*-orf477Δ-*Tn2* transposon unit (TU) (Supplementary Figure 1), similar to integration sites described previously (28, 29). This TU was transposed into the 1,617 bp *mppA* gene (encoding murein peptide permease A), which was split into 327 bp and 1290 bp segments (at NCTC13441 genome coordinates 2,522,100-2,523,713 bp). No direct repeats flanking the *bla_CTX-M-15_* element were observed. The *bla_CTX-M-15_* was separated upstream by a 48 bp spacer sequence from a fragmented ISEcp*1* upstream adjacent to IS26, and downstream *bla_CTX-M-15_*was separated by 46 bp spacer from an orf477 segment, which was flanked by an incomplete Tn*2* and IS26 elements at the 3’ and 5’ ends (Supplementary Table 2, Supplementary Figure 1), suggestive of one-ended transposition or a deletion following transposition (28, 29). The fourth assembly from C2_7 (ERR191697) contained a *bla_CTX-M-15_* gene on an IncFII/FIA plasmid with an incomplete Tn*2* element and a fragmented ISEcp*1* (p_*bla_CTX_*_-M-15_-orf477Δ-Tn*2*) flanked by IS*26* elements (Supplementary Table 2, Supplementary Figure 1). The fifth assembly was from C1 (ERR191724) and had a *bla_CTX-M-14_*-positive pV130-like IncFII plasmid (100% identity) with an intact ISEcp*1* at the 5’ end and an incomplete copy of IS*903B* at the 3’ end (p_*ISEcp1-bla_CTX_*_-M-14_-IS*903B*) (Supplementary Table 2, Supplementary Figure 1).

### Genomic context of blaCTX-M-15 the Irish collection highlight genetically diverse C subclades

Our findings indicated that the chromosomal *bla_CTX_*_-M-15_ TU inserted into the chromosome was a potentially unique characteristic of the Irish LTCF C2_8 isolates, in contrast to the plasmid-associated *bla_CTX-M-15_*in other C2 isolates, and plasmid-associated *bla_CTX-M-14_* in C1 identified by the PacBio sequencing (Supplementary Figure 1). This was tested in 54 Clade C isolates from the Irish LTCF by resolving the exact genomic architecture of regions with the *bla_CTX-M_* by genome assembly and mapping reads to construct a phylogeny (Supplementary Figure 2). Assemblies of the 54 were compared with the PacBio references and NCTC13441 (ERR718783) and the *bla_CTX-M_*, IS*Ecp1*, Tn*2*, IS*903B*, and *mppA* copy numbers were inferred from read mapping distributions, including verification of reads spanning the genetic elements and TU boundaries (Supplementary Figure 3). Of the 54, 38 were *bla_CTX-M-15_*-positive (all C2), nine were *bla_CTX-M-14_*-positive (all C1), five had no *bla_CTX-M_* gene (n=3 from C2, n=2 from C1), and two had both *bla_CTX-M-15_*and *bla_CTX-M-14_* genes (ERR191646 and ERR191657 from C2_8) (Supplementary Figure 3). C2_8 isolates (n=29) had a chromosomal insertion of *bla_CTX_*_-M-15_ (Supplementary Figure 4), contrasting with C2_7 (n=9) that typically had a fragmented IS*Ecp1* with a plasmid-associated *bla_CTX_*_-M-15_ gene like the C2_8 and C2_7 PacBio reference strains references (Supplementary Figure 5). The C2_9 (n=5) isolates had a plasmid-bound *bla_CTX_*_-M-15_ gene adjacent to a 496 bp IS*Ecp1* fragment (p_shortIS*Ecp1*-*bla_CTX_*_-M-15_-orf477Δ-Tn*2*, Supplementary Figure 5). Like the PacBio C1 assembly above, the C1 (n=11) isolates had a plasmid-associated IS*Ecp1-bla_CTX_*_-M-14_-IS*903B* TU with three IS*Ecp1* copies along with a duplicated *bla_CTX_*_-M-14_ gene, though two were *bla_CTX_*_-M_-negative.

Examining the rest of the collection in the same way showed that the *mppA* TU insertion was unique to the 41 Irish LTCF isolates in Clade C2_8 and this mutation was not found among any of the other 63 isolates from Ireland either in LTCF, community or hospitals. This is consistent with a pattern of clonal expansion in the LTCF. Of the 690 global isolates, 11 of the 19 with a disrupted *mppA* gene were *bla_CTX-M-15_*-positive and clustered within the clonally expanded C2_8 *mppA-*insertion lineage. The remaining eight were independent events: six had no *bla_CTX-M_* gene and one had a *bla_CTX-M-19_* gene. Across all 794, C2 had a high rate of *bla_CTX_*_-M-15_-positives isolates, reiterating the correlation of *bla_CTX_*_-M-15_ with the expansion of C2, with incidences of 84% in C2_7 isolates (303 out of 362), 83% in C2_8 (71 out of 86), and 67% in C2_9 (76 out of 113).

### Time of origin of the ST131 clones

The estimated time of the most recent common ancestor (TMRCA) of different phylogenetic groups was investigated with BEAST. We estimated a mutation rate of 4.14×10^-7^ SNPs/site/year (95% highest posterior density [HPD] intervals 3.74-4.57×10^-7^), equivalent to 1.858 mutations/genome/year. A dated phylogeny (Figure 4) of all 794 isolates estimated a TMRCA for ST131 of around 1901 (95% HPD intervals 1842-1948). Clade C originated in 1985 (95% HPD 1980-1989). The FQ-R C1/C2 ancestor originated in 1992 (95% HPD 1989-1994), more recently than previous estimates of 1987 (12) and 1986 (15). Following this event, C1 and C2 diversified in parallel around 1994 (95% HPD 1991-1996, 95% HPD 1992-1995, respectively). C2 is composed of divergent subclades C2_7, C2_8 and C2_9. C2_7 diversified from the C2_8/9 lineage in 1995 (95% HPD 1993-1997). Finally, a group of strains radiated within C2_8 and formed a “displacement clade” (Figure 5A).

**Figure 4.**
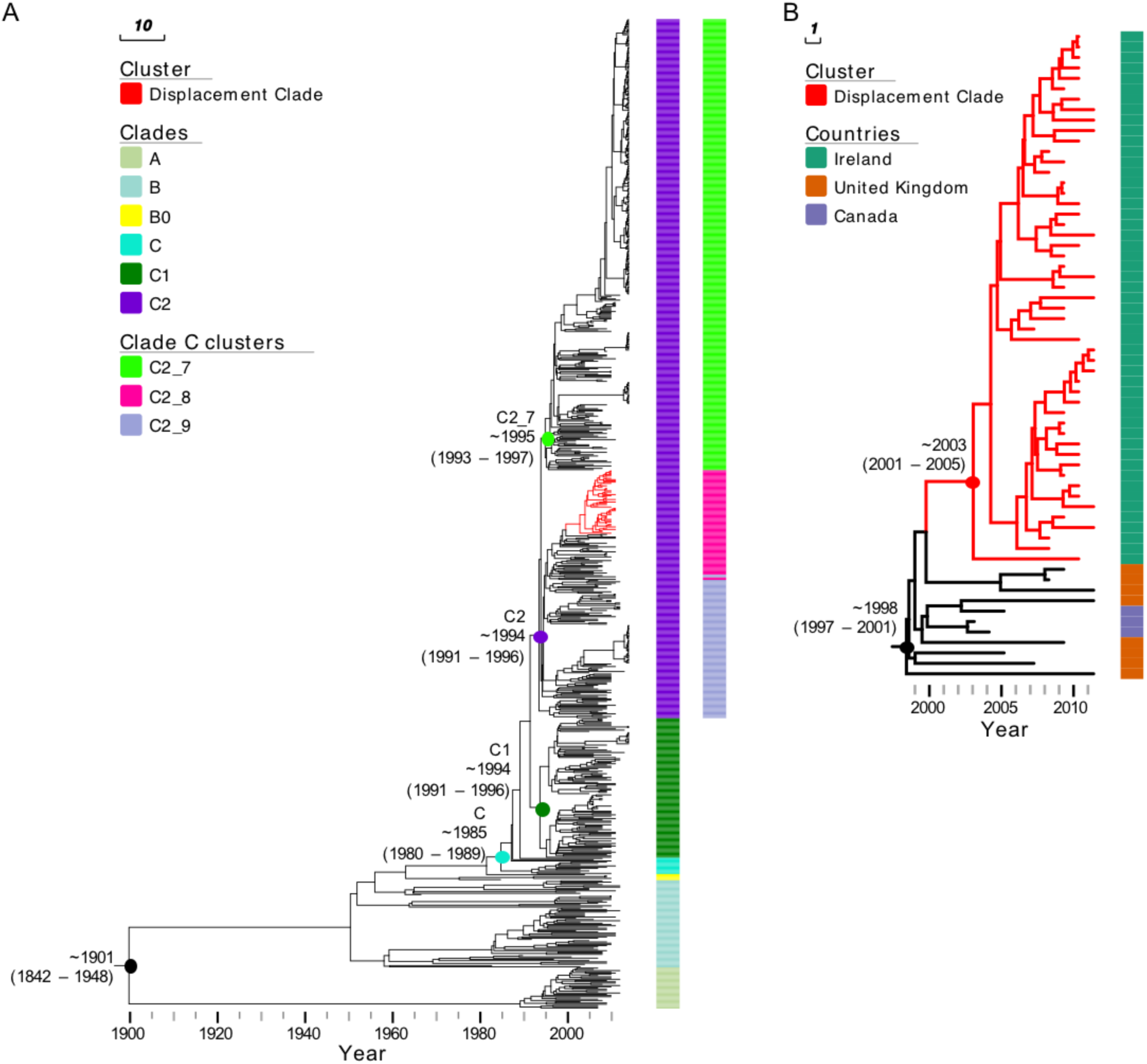
Bayesian maximum clade credibility tree of *E. coli* ST131 isolates. (A) Phylogeny of 794 isolates analysed in this study. The tree is annotated with column representing major phylogenetic clades (Clades) as well as subclades within clade C (clade C clusters). The estimated TMRCA for major clades is shown on the tree. Branches of the cluster representing isolates from the Irish LTCF displacement clone are coloured in red. (B) A higher resolution view of the Irish LTCF displacement clone, annotated with colour strips representing isolate’s country of origin.

The TMRCA of the C2_8 clade was estimated at around 2003 (95% HPD 2001-2005) and all isolates in this clade contained a chromosomal *bla_CTX-M-15_* inserted between a truncated *mppA* gene. The “displacement clade” within the C2_8 subcluster comprised of 41 Irish LTCF isolates from the local collection which had a unique TU insertion in the *mppA.* This is in addition to ten other Irish isolates from clinical or community sources (n=51 in total) with a mutant *mppA* but showed a different TU insertion. The 11 *bla_CTX-M-15_*-positive isolates that clustered with the Irish C2_8 isolates also had a disrupted *mppA* gene and were from the UK (n=8) and Canada (n=3). Together, these 62 shared a TMRCA of around 1998 (95% HPD 1997-2001), indicating that the *mppA* insertion may have occurred in the ancestral branch dating to 1996-1998 in the UK or North America (Figure 5B).

This evidence highlighted a single genetic origin of the ancestral C2_8 lineage in the Irish LTCF (Figure 4), though it was rare until 2009 (Table 3), potentially presenting opportunities for multiple introductions of C2_8. Prior to 2008, C1_10 was most common, consistent with a pattern of replacement by C2_8 with the mutant *mppA* insertion that clonally expanded. Nine out of 12 isolates from this facility detected between 2005-2007 belonged to C1_10. This was the group of isolates corresponding with the outbreak identified in 2006. Conversely, in the 57 samples isolated from 2008-2011, all were *bla_CTX-M-15_* positive, and 36 were classified in C2_8 (four of which were also both *bla_CTX-M-14_*-positive), seven in C2_9 and eight in C2_7. In the global isolates, in contrast to the LTCF, C2_8 accounted for only 5% of isolates, whereas C1_10 and C2_7 accounted for 14% and 49% (respectively) with no evidence of this clonal displacement outside of the LTCF.

**Table 3.**
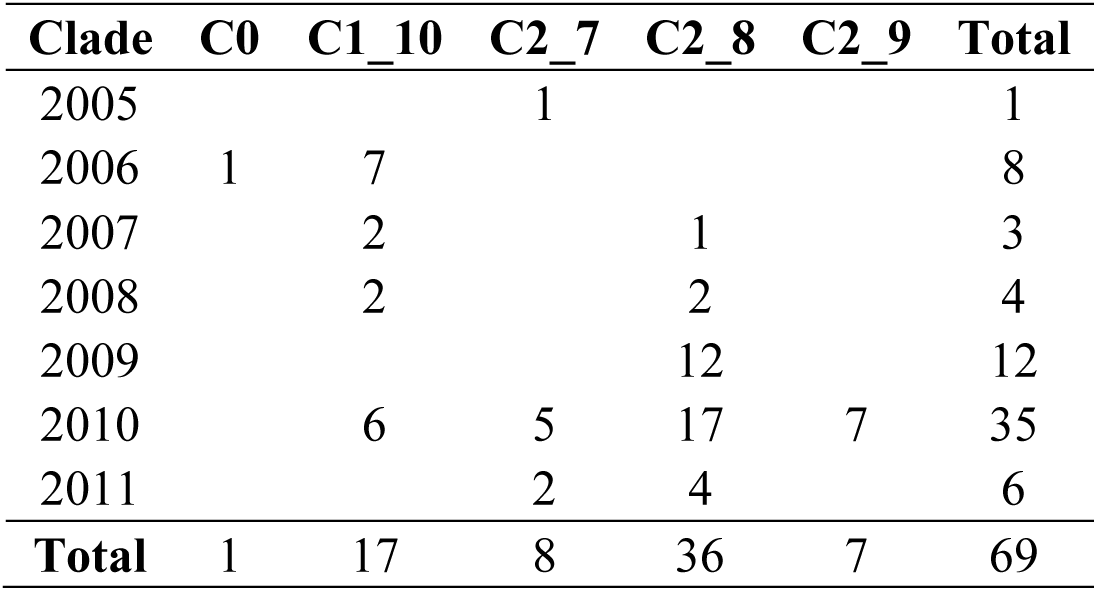
The numbers of isolates from the LTCF in Ireland (n=69) across the ST131 clades showed that C1_10 was most common at the outset of the study, and that C2_8 became more prevalent after 2008, suggesting a possible replacement and clonal expansion of this lineage.

## Discussion

Here, we traced the genomic background of ESBL-*E. coli* ST131 isolates collected from residents of a LTCF in Ireland where an outbreak was recognised in 2006. The relationship between the isolates was first identified based on indistinguishable pulsed field gel electrophoresis (PFGE) patterns among 18 patients (26). Since the outbreak was detected in 2006, there has been extensive progress in the higher discriminatory power of genome-sequencing compared to PFGE and other typing tools, such as MLST (30–32). To gain a further understanding of the origins of the outbreak and to observe changes in *E. coli* population structure in LTCF residents, we performed whole genome sequencing of all ST131 ESBL-*E. coli* isolates submitted from the LTCF over seven years. We compared these to 35 other ST131 isolated in Ireland; 9 from other LTCFs, 2 from the community and 24 from hospitals including 14 from the referral hospital and 10 from 3 other hospitals, in addition to 690 ST131 from global datasets.

We identified distinct genetic clusters within this set of 794 closely related isolates based on core genome phylogenetic signals, and as in previous studies (11), we identified subclade C2 as the most abundant ST131 group accounting for 71% of the entire collection. Four genetic subgroups were common in the specific LTCF, one from subclade C1 (C1_10) and three from C2 (C2_7, C2_8, C2_9). The resident ST131 lineage (C1_10) in the LTCF in the period 2005-2007 was the cause of the initial outbreak investigation, but surprisingly a newly introduced ST131 variant (C2_8) was much more common by 2009, indicating displacement of *bla_CTX-M-14_*-positive C1 isolates and clonal expansion by a genetically distinct *bla_CTX-M-15_*-positive C2 lineage within the LTCF. This pattern of clonal displacement has not yet been published for *E. coli,* but is common in other species such as methicillin-resistant *Staphylococcus aureus* (MRSA) where it can be driven by inter-hospital transfer of patients (33).

In this study, we analysed the largest global collection of whole genome data on ST131 *E. coli* and estimated the emergence of ST131 in approximately 1901. The clonal expansion of C2 in 1994 identified here was similar to Kallonen et al (15) and Zakour et al (12), who reported 1990 and 1987, respectively. We dated the C2_8 LTCF lineage to have emerged in 2001-2005 and we postulate that the clone originated in the UK or North America in 1996-1998. This was consistent with the first observation of *bla*_CTX-M-15_-positive cephalosporin-resistant *E. coli* isolated in 2001 in three locations in Britain and Northern Ireland (34, 35). However, C2_8 was generally not as successful as C2_7, which emerged around the same time (1995) and disseminated globally. It has been suggested that the evolution of C2 subclades has been shaped by the acquisition of IncFII plasmids encoding *bla_CTX-M-15_* (11), which was also observed here for C2_7. We extend this by showing that *bla_CTX-M-15_* in C2_8 was mobilized from IncFII plasmids by IS*Ecp1*-mediated transposition to the chromosome at *mppA* in a TU structured as IS*Ecp1*-*bla_CTX-M-15_*-orf477Δ-Tn*2*. The high copy number and fragmented pattern of IS*Ecp1,* which enabled a chromosomal insertion, was found for different *bla_CTX-M_* alleles in *E. coli* and may be linked to altered expression of the gene on the chromosome relative the plasmid (13, 14). Our work shows although ST131 is disseminated globally, evolutionary events have resulted in the clonal expansion of new lineages, such as C2_7 globally and C2_8 locally in one LTCF. This has coincided not only with the horizontal gene transfer of plasmids encoding *bla_CTX-M-15_*or *bla_CTX-M-14_*, but also the chromosomal insertions like *bla*_CTX-M-15_ in C2_8 followed by vertical transmission, and also *bla*_CTX-M-14_ 5’ of the chromosomal *rlmL* gene in one C1 isolate (ERR191666).

In conclusion, we investigated an outbreak of ESBL *E. coli* ST131 in a LTCF in Ireland and observed changes in this LTCF different to the global pattern. We found that the outbreak began with a Clade C1 strain encoding *bla_CTX-M-14_* gene on a plasmid, and that this lineage was displaced by a Clade C2 strain with a chromosomally-encoded *bla_CTX-M-15_* gene. Both lineages associated with the LTCF are resistant to broad-spectrum cephalosporins and the selective forces in this specific niche driving lineage displacement are unclear.

This highlighted the importance of long-read sequencing to resolve plasmids and to decipher plasmid and chromosomal spread of ESBL genes. The ability of long-read sequencing to identify novel plasmids and extra-chromosomal elements such as bacteriophage that do not integrate but replicate chromosomally, should become a new standard. The sustained use of ciprofloxacin and third generation cephalosporins will continue to enrich for Clade C2 lineages and mobile genetic elements, highlighting the global need to reduce the selective pressure from these antimicrobials. The diversity of ST131 lineages and resistance elements indicates a need for surveillance strategies to identify ST131 subclones, plasmids and transposable elements. The characterisation of those specific properties that make specific lineages successful in particular contexts remains one of the key challenges in understanding the dynamics of emergence and spread of new variants of common bacterial species. Focused attention to successful strains could help to explore these interactions and control the epidemic of *E. coli* resistance.

## Materials and Methods

### Irish bacterial isolate collection and short read genome sequencing

A total of 90 *E. coli* ST131 isolates from Ireland were isolated and sequenced. Among these 90, 69 were sampled from 63 residents during 2005-2011 from a single LTCF with an outbreak of ESBL-producing *E. coli* in 2006 (26) and 21 were clinical isolates from the referral hospital (Galway University Hospital: n=8 hospitalized patients, n=11 residents of other Irish LTCFs, and n=2 community isolates submitted from general practitioners).

Bacterial genomic DNA for the 90 isolates was extracted using the QIAxtractor (Qiagen, Valencia, CA, USA) according to the manufacturer’s instructions. Library preparation was conducted according to the Illumina protocol and sequenced (96-plex) on an Illumina HiSeq 2000 platform (Illumina, San Diego, CA, USA) using 100 bp paired-end reads. On average, 5,014,175 (range 3,489,126-8,166,084) raw sequence reads were generated per isolate, with a mean insert size of 260 (range 244-280).

### Complementary datasets

For context, DNA read libraries and associated metadata were retrieved for 704 *E. coli* ST131 isolates, 14 of which were BSIs from four referral hospitals in Ireland and 4/14 isolates were obtained from the referral hospital (Galway University Hospital). The remaining global 690 were isolated between 1967 and 2014 and included 167 (clinical=155, environmental=7, unknown=5) isolates obtained from global collections (16, 37), 297 from a UK LTCF (22), and 226 were associated with BSI in the UK (15, 22) (Supplementary Table 1).

### Long read sequencing, assembly and annotation

DNA was extracted using the phenol-chloroform method (27) and sequenced using a PacBio RSII Instrument (Pacific Biosciences, Menlo Park, CA, USA) for five isolates (ERR191646, ERR191657, ERR191663, ERR191724 and ERR191697). Sequence reads were assembled using HGAP v3 (38) of the SMRT analysis software v2.3.0 (https://github.com/PacificBiosciences/SMRT-Analysis), circularized using Circlator v1.1.3 (39) and Minimus 2 (40), and polished using the PacBio RS_Resequencing protocol and Quiver v1 (https://github.com/PacificBiosciences/SMRT-Analysis). This assembled the plasmids for each of the isolates used as references for short read mapping. NCTC13441’s HDF5 files were converted to FASTQ with 308,854 reads using pbh5 tools (smrtanalysis v2.3.0p4). These reads were screened for PacBio adapter sequence using Cutadapt v1.9.1 and corrected using BayesHammer from SPAdes v3.0.0 with a seed k-mer of 127, yielding a total of 41,813 reads.

### Genome assembly, read mapping, AMR gene identification and plasmid typing of the 794

*De novo* assembly of short read data for the 794 libraries was performed using VelvetOptimiser v2.2.5 (41) and Velvet v1.2 (17). An assembly improvement step was applied to the assembly with the best N50, whose contigs were scaffolded using SSPACE (42) and contig gaps reduced using GapFiller (43). The assembly pipeline generated an average total length of 5,166,846 bp (range 4,697,700-5,460,279 bp) from 97 contigs (range 31-486) with an average contig length of 59,340 bp (range 11,186-1,661,401 bp) and an N50 of 227,849 bp (range 30,788-763,538 bp) (Supplementary Table 3). Assemblies annotated using Prokka v1.5 (44) and a genus-specific database from RefSeq (45). The 794 short read libraries were mapped to NCTC13441 genome (accession ERS530440) (22), PacBio assemblies and reference plasmids using SMALT v7.6 (http://www.sanger.ac.uk/resources/software/smalt/). The genomic locations of the *bla*_CTX-M_ genes and nearby MGEs were examined by aligning the short and long read assemblies using BLAST to the *bla*_CTX-M_-positive TU isoforms, including one with a split *mppA* gene containing the TU (Supplementary Figure 3). The two observed *mppA* isoforms were recorded as T for truncated (separated 327 bp and 1290 bp segments) or I for intact (Supplementary Table 1). SNP screening at *mppA* across the 794 showed limited variation: just one doubleton and four singleton SNPs.

AMR genes in the 794 were identified by alignment with the 2,158 gene homolog subset of the Comprehensive Antibiotic Resistance Database (CARD) v1.1.5. Plasmid incompatibility group and replicon types were identified (Supplementary Table 4) by comparing the genomes against the PlasmidFinder database (accessed date 16/03/17) (46) with a 95% identity threshold.

### Quality control, genome assembly and read mapping of 54 Irish read libraries

Adapter sequences in the libraries of the 54 Irish Clade C reads were trimmed with Trimmomatic v0.36 (47) using a Phred score threshold of 30 (Q30), a ten bp sliding window and a minimum read length of 50 bp. On average, these had 2,400,763 reads initially, of which 7.8% were removed by trimming. These were corrected using BayesHammer in SPAdes v3.9. The effects of removing low-quality bases and reads was quantified using FastQC v0.11.5 with MultiQC v1.3, which showed base-correction removed an additional 14.3% of reads on average, leaving a mean of 1,898,990 per library. This showed levels of base quality and potential contaminants were consistent across the libraries.

Read libraries of the Irish 54 were assembled into contigs using SPAdes v3.9 with a k-mer of 77 (48). This optimal k-mer maximised the N50 value determined by Quast v5.0 (49). The contigs were ordered and scaffolded based on the NCTC13441 reference chromosome, plasmid and annotation using ProgressiveMauve (50), producing an average scaffold N50 of 177,758±12,199 (mean±SD) bp with a mean assembly length of 5,434,674±153,210 bp and an average of 234 contigs per library.

A total of 59,536 bases at low complexity repeats, homopolymers, sites within 1 Kb of chromosome edges, bases within 100 bp of a contig edge, or at tandem repeats were masked from the NCTC13441 reference chromosome using Tantan v0.13 (www.cbrc.jp/tantan/), which was indexed using SMALT v7.6 using a k-mer of 19 with a skip of one, as were all reference sequences here. The short read libraries were mapped to reference sequences using SMALT v7.6, and the resulting SAM files were converted to BAM format, sorted and PCR duplicates removed using SAMtools v1.19. The MGE, *mppA* and *bla*_CTX-M_ gene structures were examined by alignment as above so that local copy number changes, mapping breakpoints and read pileups could be screened by mapping Illumina reads to the PacBio and contig references. The local gene structure was visualised with R v3.5.2 and the MARA Galileo AMR database (51, 52).

### Phylogenetic analysis of 794 isolates

To construct phylogenies reflecting the genealogical relationships and evolutionary changes, SNPs were identified using Gubbins v2.3.4. The SNPs arising by mutation were used to create a maximum-likelihood midpoint-rooted phylogeny using RAxML v8.0.19 (53) using a General Time Reversible + gamma (GTR+G) substitution model with 100 bootstraps across 362,009 sites. Phylogenetic trees were visualised with iToL (http://itol.embl.de) (54) and FigTree v1.4.3 (http://tree.bio.ed.ac.uk/software/figtree/) (55). For the 54 Irish Clade C collection, a phylogeny was created as above with RAxML with 100 bootstraps, and a network was constructed using uncorrected p-distances with Splitstree v4.14.2 (56), visualized with FigTree.

### Inference of subclade common ancestry and historical population size changes

To reconstruct time-calibrated phylogeny ST131 we used a core genome alignment of 794 isolates that contained 8,567 SNPs after the exclusion of regions representing MGEs, recombinant tracts and sites with an uncalled genotype across >1% of sequences. Each sequence in the alignment was annotated with the year of isolation. The strength of the molecular clock signal was measured by linear regression of the root-to-tip genetic distance against year of sampling using TempEst (55), which revealed a correlation coefficient of R^2^ = 0.4. Bayesian inference of phylogeny was performed with BEAST v2.4.7 (57) based on a GTR+G nucleotide substitution model. To optimise computing efficiency in a large dataset, model selection was implemented on a subset of isolates (n=205) that tested two clock rates (strict versus relaxed uncorrelated lognormal) across three population models (constant, exponential and Bayesian skyline). Five replicates for each of the six models were tested. The MCMC chain was run for 50 million generations, sampling every 1,000 states. Log files from the five independent runs per model option were assessed for convergence using Tracer v1.5, and combined after removal of the burn-in (10% of samples) using LogCombiner. The relaxed lognormal clock with Bayesian skyline model was the best fit, consistent with previous work (58) and so this was used to model the evolutionary history across all 794 isolates with 15 replicates. The maximum clade credibility (MMC) tree was generated with TreeAnnotator.

## Acknowledgments

This project was funded by a Dublin City University (DCU) O’Hare Ph.D. fellowship, a DCU Enhancing Performance grant, and the Health Innovation Challenge Fund (WT098600, HICF-T5-342), a parallel funding partnership between the Department of Health and Wellcome Trust. Catherine Ludden was a Wellcome Trust Sir Henry Wellcome postdoctoral fellow (110243/Z/15/Z). Derek Pickard was supported by funding from the NIMR Cambridge BRC AMR theme and the ESPRC Vaccine Hub.

The views expressed in this publication are those of the authors and not necessarily those of the Department of Health or Wellcome Trust. J.P. and S.J.P are paid consultants to Next Gen Diagnostics LLC.

Author contributions were as follows: conceptualization, C.L., D.M., D.P., J.P., M.C. and S.J.P.; methodology, A.G.D., C.L., D.J., and T.D.; formal analysis, A.G.D., C.L., D.J., and T.D.; investigation, A.G.D., C.L. and T.D.; visualization, A.G.D., C.L., D.J., and T.D..; writing of original draft, A.G.D., C.L., D.J., J.P, and T.D.; writing, reviewing, and editing, A.G.D., C.L., D.J., J.P., M.C., S.J.P. and T.D.; funding acquisition, C.L., D.M., J.P., M.C. and S.J.P.

**Table 1. Distribution of fimH alleles across the entire collection of ST131 (n=794).** The collection consisted of three main clades subdivided into six subclades: A (n=33), B (n=70), B0 (n=5), C0 (n=14), C1 (n=111) and C2 (n=561). The frequencies of the four most common *fimH* allele types are shown: *H41*, *H22*, *H27* and *H30*; the rest are classified as “other”. No FQ-resistance mutations were detected in *fimH22/27/41*.

**Table 2. Distribution of ribosomal sequences types (rST) and Fastbaps clusters across clade C (n=686).** The entire ST131 set (n=794) was largely composed of isolates from clade C (n=686, 86% of total) that was categorised into five subclades by Fastbaps clustering: C0 (n=14, clusters 2-5 and 11), C1 (n=111, cluster 10), C2_7 (n=362, cluster 7), C2_8 (n=86, cluster 8) and C2_9 (n=113, cluster 9). The National (n=104) and global (n=690) ST131 had two main ribosomal sequence types (rSTs): rST1850 associated with the Irish C2_8 LTCF set (85%), and rST1503 that often corresponded to C2_7 (92.5%). Fastbaps clusters 2, 3, 4 and 5 in C0 represented one isolate each – only cluster 3 was *bla_CTX_*_-M-15_-positive.

**Supplementary table 1.** Isolate information and metadata of the 794 global ST131 collection.

**Supplementary table 2.** PacBio sequence annotation of regions encoding *bla_CTX-M-15_* and *bla_CTX-M-14._* in the IncFII/FIA *bla_CTX-M-15_* reference plasmid pEK499 (GenBank: EU935739.1), IncFII/FIA Clade C2_7 plasmid (assembled from ERR191697) and IncFII *bla_CTX-M-14_* Clade C1 plasmid (assembled from ERR191724). Annotations were created using the Galileo AMR online tool. **\***Arrows indicate the orientation of features. **^#^**In text annotations indicates that an incomplete copy of the feature is present. The dashed part of arrow indicates which end is missing. Annotations of partial features may not distinguish correctly between particular variants.

**Supplementary table 3.** Assembly metrics for the 794 global ST131 collection.

**Supplementary table 4.** Plasmid replicon typing for the 794 global ST131 collection.

---

**Supplementary Figure 1.**
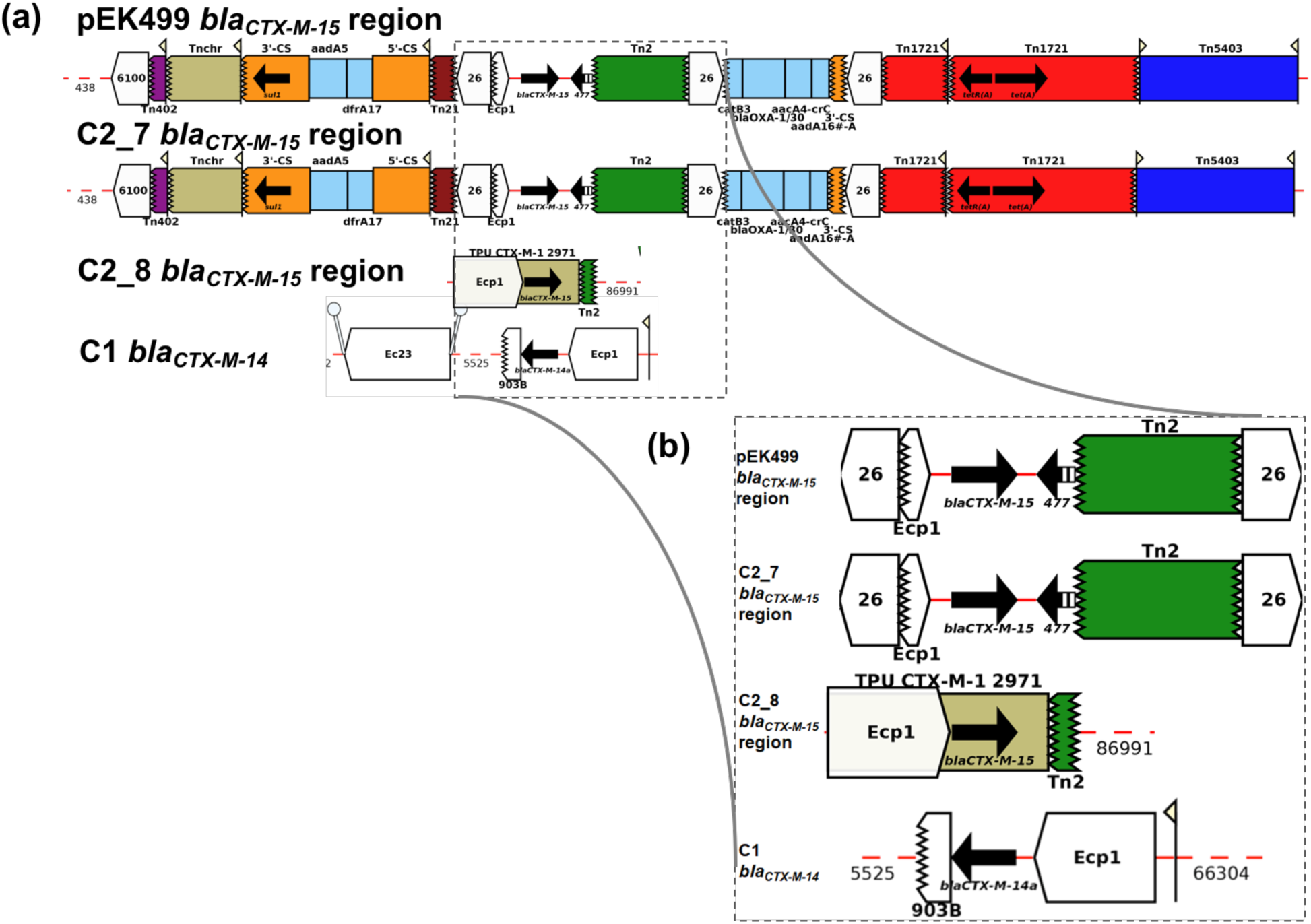
Flanking genetic elements of blaCTX-M genes in pEK499 and in the chromosome or plasmid-derived contig of C2_7, C2_8 and C1 strains. A blaCTX-M-15 element on the IncFII/FIA pEK499 reference and in subclade C2_7, a chromosomal blaCTX-M-15 gene in C2_8 isolates and a blaCTX-M-14 element on IncFII subclade C1 plasmid (a). The region encoding blaCTX-M-14 and blaCTX-M-15 are highlighted in a black box. Arrows indicate the orientation of features, with the forward direction defined as the direction of transcription for genes, towards the main part of the attC site for cassettes, in integrons towards attI for 5’ flanking regions, away from the cassette array for 3’-flanking regions, relative to the direction of transcription of the transposase gene for insertion sequences and transposons (Tn) (ie, inverted repeat left to inverted repeat right) and to the direction of the reverse transcriptase for Group II introns. The missing end of a feature is shown by a zig-zag line. The inset represents the area bound by the dashed line and is shown in more detail in (b). Read mapping copy number for n=27 C2_8 isolates showing that all had 1+ IS*Ecp1* elements (blue) 5’ of a *bla_CTX-M-15_* gene (red) with consistent coverage spanning the *mppA* gene (green) including reads spanning each elements’ breakpoints, with the TU isoform chr_shortIS*Ecp1-bla_CTX_*_-M-15_-shortTn*2*. N=13 (a) had IS*Ecp1* fragments of 1,203 bp, 529 bp, 76 bp and 76 bp with one *bla_CTX-M-15_* gene, though within this some had one or two extra 76 bp IS*Ecp1* fragments on their plasmids, including 8289_1#24 (ERR191657 in Table 3, not shown here) and 8289_1#53 (not shown here). 8289_1#24 had a *bla_CTX-M-14_* gene, and had 74-77 bp IS*Ecp1* fragments on its plasmid, as well as 76 bp, 529 bp and 1203 bp IS*Ecp1* segments on its chromosome. 8289_1#5 (ERR191638) also had its *bla_CTX-M-15_* gene was 57,725 bp distant from the IS*Ecp1* fragments. 8289_1#38 (ERR191671), 8289_1#61 (ERR191694) and 8289_1#95 (ERR191728) had a 76 bp IS*Ecp1* fragment adjacent to the chromosomal *bla_CTX-M-15_* gene at 27,742 bp (8289_1#38), 6,835 bp (8289_1#61) and 32,815 (8289_1#95) from the other IS*Ecp1* copies, consistent with recombination between IS*Ecp1* segments. N=7 (b) were like this previous group, but without the 529 bp IS*Ecp1* fragment. Another set (n=5, third panel) had a full 1,655 bp IS*Ecp1* element with no fragmentation. One isolate (8289_1#54) had three full 1,655 bp IS*Ecp1* elements on both its chromosome and plasmid, and fragments of 76 bp and 74 bp adjacent to the *bla_CTX-M-15_* gene (c). Finally, the strain ERR191729 (8289_1#96) had three *bla_CTX-M-15_* gene copies and a first IS*Ecp1* fragment 529 bp where the TU was inverted and duplicated, and separate from a 1,203 bp IS*Ecp1* at mppA, suggesting that recombination between the chromosomal and plasmid IS*Ecp1* IRs may have transferred the *bla_CTX-M-15_* gene back to the plasmid (e). The tables (right) shows the IS*Ecp1* assembly coordinates, spacer DNA lengths and *bla_CTX-M-14_* assembly coordinates.

**Supplementary Figure 2.**
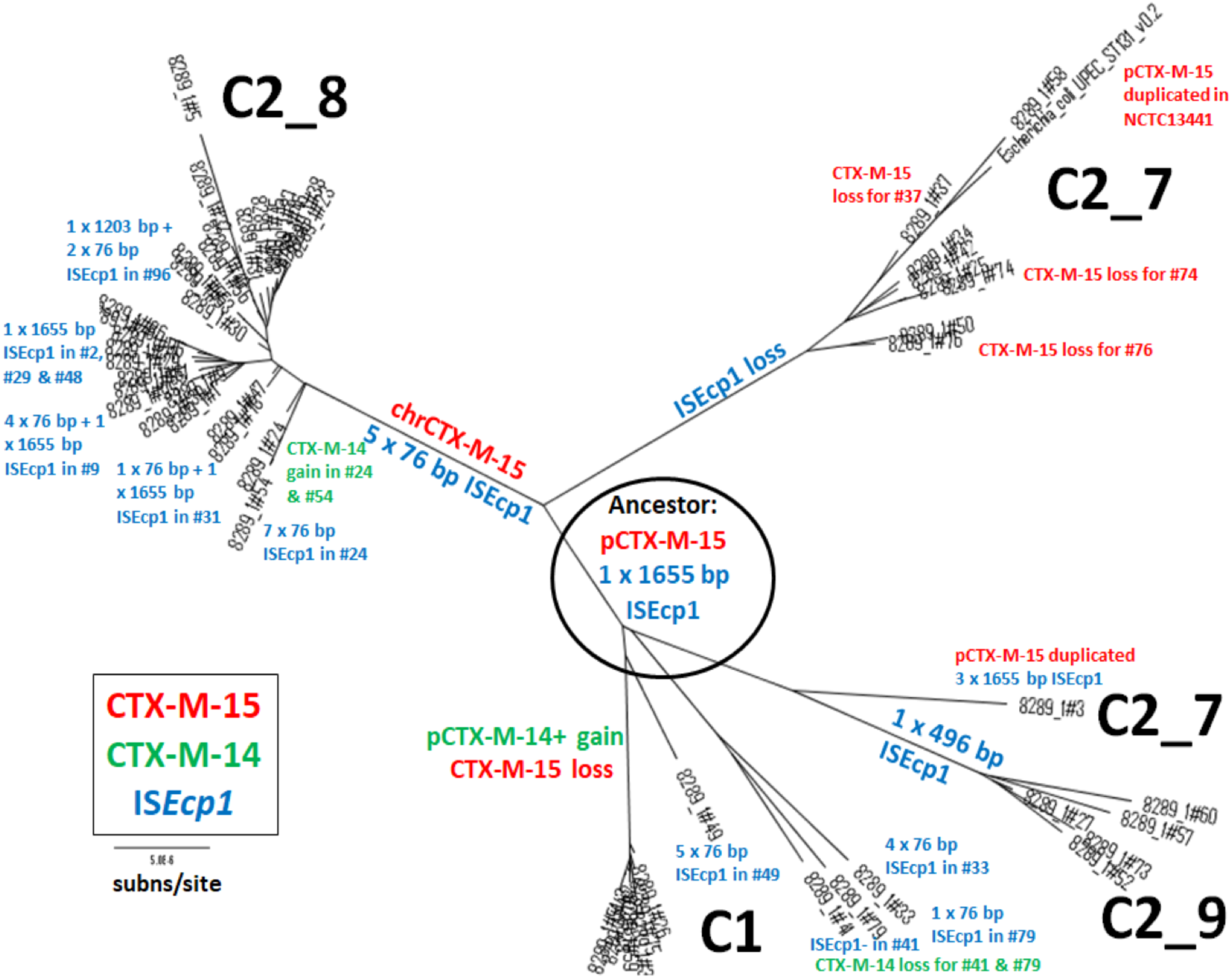
Phylogenetic reconstruction of N=54 ST131 strains collected from Irish long-term care facilities. A phylogenomic network of the 54 Irish Clade C samples’ chromosomal mutational SNPs built using RAxML and ClonalframeML and drawn with FigTree v.1.4.3. The phylogeny of the n=54 was rooted using the topology of n=794, which C1 as the most divergent lineage, with C2_9 diverging next, followed by C2_8 and C2_7, though here the smaller sample size meant that the ancestral lineage was unclear and so could be approximated by the C1-C2 origin, where the ancestor likely had a plasmid with a 1,655 bp IS*Ecp1* 5’ of a *bla_CTX-M-15_* gene. The *bla_CTX-M-15_* gene changes are in red, the *bla_CTX-M-14_* gene mutations are in green, and the IS*Ecp1* differences are in blue. The subclades C1, C2_7, C2_8 and C2_9 are shown, and the scale bar shows five substitutions per Mb.

**Supplementary Figure 3.**
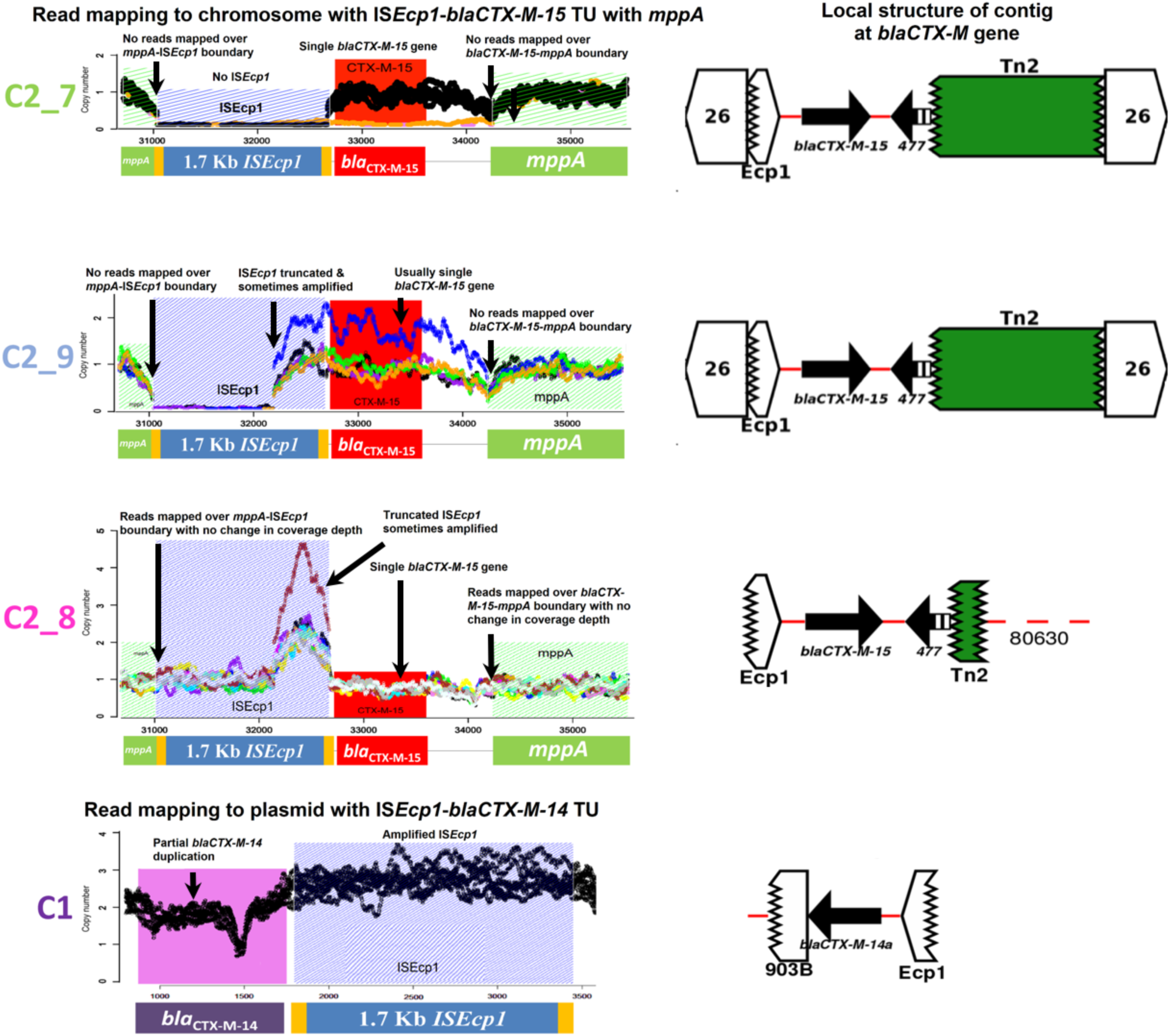
Genomic context of *bla*_CTX-M_ genes in C2_7, C2_9, C2_8 and C1 isolates. Read mapping copy numbers for 33 of the 54 the Irish isolates from C2_7 (n=6 shown), C2_9 (all n=5 shown), C2_8 (n=15 shown) and C1 (n=7 shown) across IS*Ecp1* elements (blue), the *bla_CTX-M-15_* gene (red), the *bla_CTX-M-14_* gene (pink) or the *mppA* gene (green). C2_8 all had consistent coverage of the chromosomally inserted TU isoform IS*Ecp1-bla_CTX_*_-M-15_-shortTn*2* spanning *mppA* and typically had with IS*Ecp1* fragments of 1,203 bp, 529 bp, 76 bp and 76 bp. The C2_9 isolates had a 496 bp IS*Ecp1* element and a *bla_CTX-M-15_* gene. Most C2_7 isolates had no IS*Ecp1* and one *bla_CTX-M-15_* gene. C1 had the TU isoform p_IS*Ecp1-bla_CTX_*_-M-14_-IS*903B* with a duplicated IS*Ecp1* element and duplicated *bla_CTX-M-14_* gene. Reads from non-C2_8 libraries mapped at *mppA* in the TU isoform, but with gaps indicating no contiguous mapping to the *bla_CTX-M-15_* gene.

**Supplementary Figure 4.**
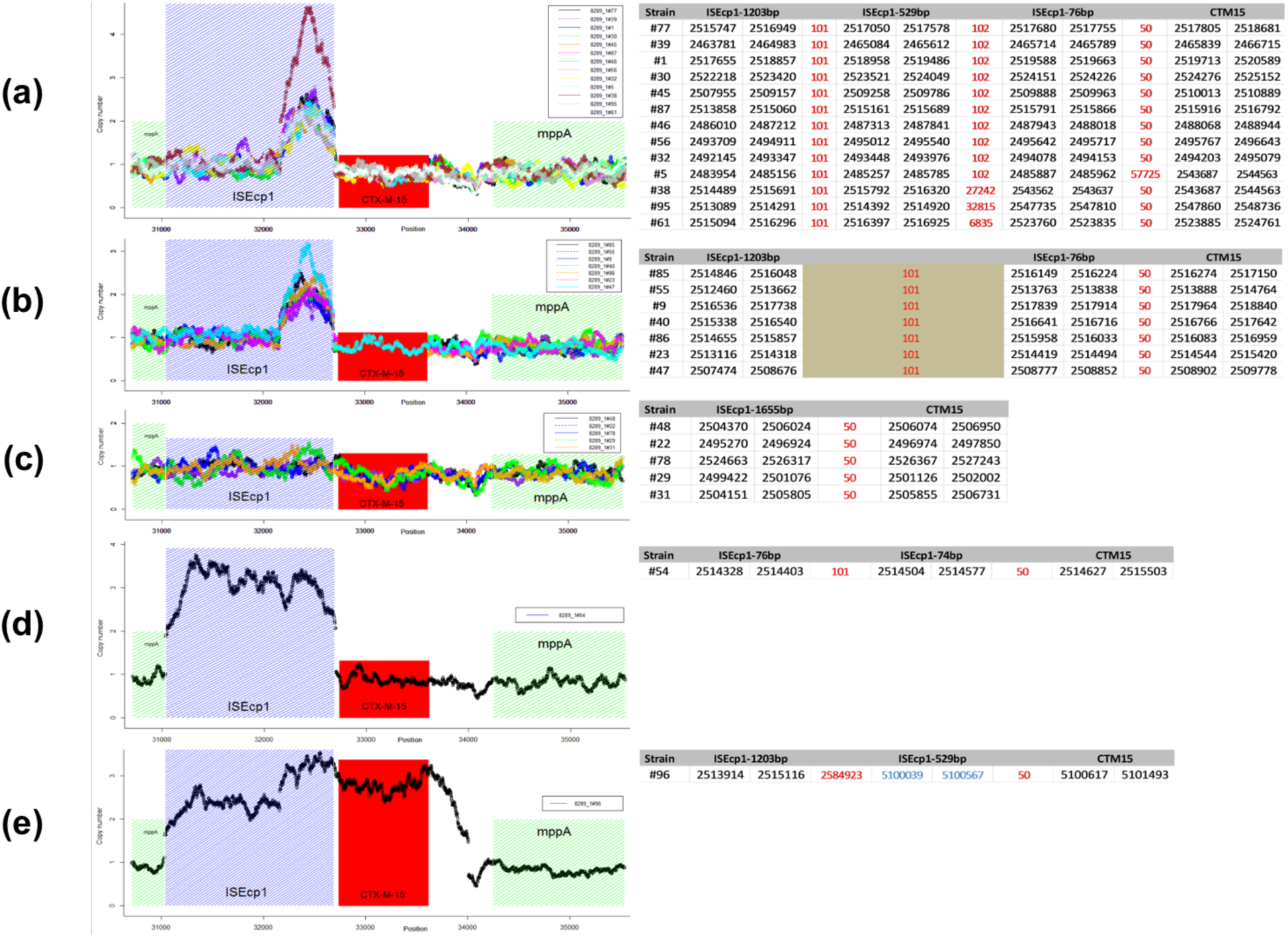
C2_8 isolates with a chromosomal IS*Ecp1-bla_CTX_*_-M-15_ at the *mppA* gene.

**Supplementary Figure 5.**
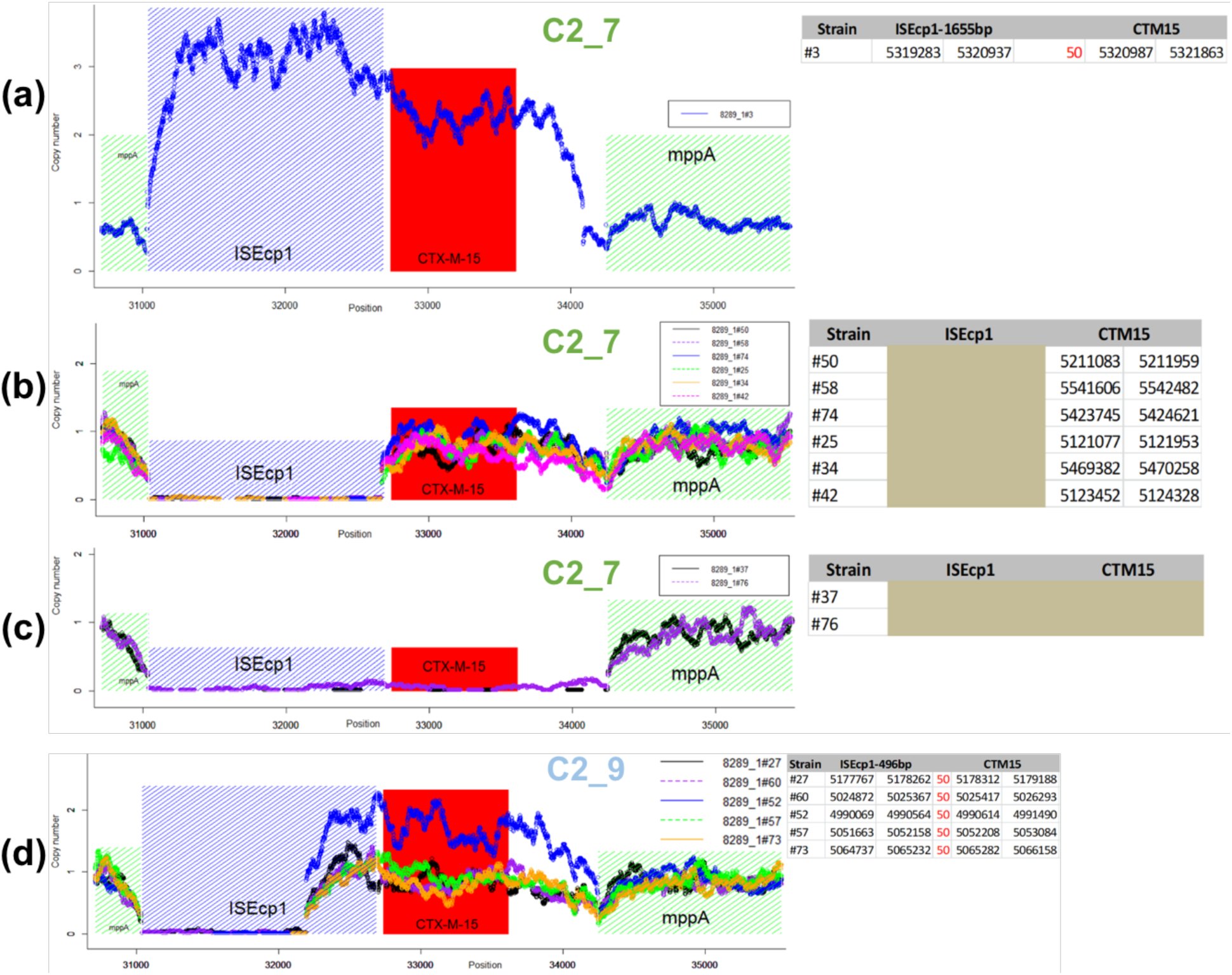
C2_7 and C2_9 strains with a plasmid-bound IS*Ecp1-bla_CTX_*_-M-15._ Read mapping copy number for n=9 C2_7 and n=5 C2_9 isolates showing one from C2_7 (8289_1#3, ERR191636, (a)) had a three IS*Ecp1* copies (blue) and a duplicated *bla_CTX-M-15_* gene (red), but no contiguity if reads mapping across to *mppA* (green). These had the TU isoform p_*bla_CTX_*_-M-15_-orf477Δ-Tn*2*. The majority of C2_7 (n=6, (b)) had no IS*Ecp1* and one *bla_CTX-M-15_* gene. Two C2_7 isolates (c) had no IS*Ecp1* and no *bla_CTX-M-14_* gene. All n=5 C2_9 isolates (d) had a 496 bp IS*Ecp1* element (blue) 5’ of a *bla_CTX-M-15_* gene (red), but no contiguity if reads mapping across to *mppA* (green) unlike C2_8. One (8289_1#52, blue) had a partial amplification of this TU. The tables on the right show the IS*Ecp1* assembly coordinates, 50 bp spacer length and *bla_CTX-M-14_* assembly coordinates.

